# Structural basis for substrate binding and specificity of a sodium/alanine symporter AgcS

**DOI:** 10.1101/293811

**Authors:** Jinming Ma, Hsiang-Ting Lei, Francis E. Reyes, Silvia Sanchez-Martinez, Maen Sarhan, Tamir Gonen

## Abstract

The amino acid, polyamine, and organocation (APC) superfamily is the second largest superfamily of membrane proteins forming secondary transporters that move a range of organic molecules across the cell membrane. Each transporter in APC superfamily is specific for a unique sub-set of substrates, even if they possess a similar structural fold. The mechanism of substrate selectivity remains, by and large, elusive. Here we report two crystal structures of an APC member from *Methanococcus maripaludis*, the alanine or glycine:cation symporter (AgcS), with L- or D-alanine bound. Structural analysis combined with site-directed mutagenesis and functional studies inform on substrate binding, specificity, and modulation of the AgcS family and reveal key structural features that allow this transporter to accommodate glycine and alanine while excluding all other amino acids. Mutation of key residues in the substrate binding site expand the selectivity to include valine and leucine. Moreover, as a transporter that binds both enantiomers of alanine, the present structures provide an unprecedented opportunity to gain insights into the mechanism of stereo-selectivity in APC transporters.

## Introduction

Controlling nutrient balance in cells and by extension entire organisms is critical. Bacteria and archaea evolved sophisticated mechanisms to adapt to life in the most extreme environments. One such example is *Methanococcus maripaludis* that exists in salt-marsh plains(1) and has the unusual ability to use both L- and D-alanine as an important nitrogen source to support life in high salt and anaerobic conditions(2–4). As a transporter that uptakes both enantiomers of alanine, the alanine or glycine:cation symporter (AgcS) from *Methanococcus maripaludis* plays essential role in this process(3). Furthermore, though L-amino acids are the dominant substrate in all kingdoms of life, it is now clear that the uptake system of D-amino acids also fulfill essential functions in many organisms. Several recent studies show that the uptake system of D-amino acids in bacteria and archaea are crucial for stationary phase cell wall remodeling(5), host metabolism and virulence(6) although the mechanism by which such amino acids are transported into cells is poorly understood.

The Amino acid-Polyamine-organoCation (APC) superfamily [Transporter Classification DataBase (TCDB)](7) represents the second largest family of secondary carriers currently known(8–11) and plays essential roles in a wide spectrum of physiological processes, transporting a large range of substrates across the membrane. Members of this superfamily have since been identified and expanded to 18 different families in diverse organisms(12). Recent structures have shown that APC superfamily members contain 10-14 transmembrane helices and exhibit sufficiently similar folds characterized by 5 or 7 transmembrane helix inverted-topology repeat motif(13). However, some of these proteins have exceptionally broad selectivity, others are restricted to just one or a few amino acids or related derivatives. It is of much interest to investigate structural basis for substrate binding and specificity of APC superfamily.

The alanine or glycine:cation symporter (AgcS) family (TC# 2.A.25) belongs to the APC superfamily and shows limited sequence similarities with other members. In contrast to ApcT and LeuT of which substrate specificity are broader(14, 15), members of the AgcS family have been reported to act as symporters, transporting L- or D-alanine or glycine with Na^+^ and/or H^+^ but no other amino acids (3, 16–20). Here, we present the crystal structure of AgcS from *Methanococcus maripaludis* in complex with L- or D-alanine as the first model for members in the AgcS family. The structure of AgcS was captured in a fully occluded conformation with the transmembrane architecture like other APC superfamily members. Functional assays demonstrate that purified AgcS binds only glycine and both enantiomers of alanine, while strictly excluding other amino acids. Further structural analyses combined with mutagenesis and biochemical studies suggest that the residues at the intracellular face of the binding pocket play a key role in substrate binding and specificity. Mutation of residues at the binding site of AgcS can expand its selectivity. Moreover, structural comparisons of AgcS with LeuT, which can only transport L-amino acids, also pave the way for a better understanding of stereo-selectivity adopted by the APC superfamily transporters.

## Results

### Overall architecture of AgcS

A *Methanococcus maripaludis* AgcS was expressed in *E. coli* and purified in various detergents for crystallization. Although successfully crystallized, AgcS alone produced only crystals diffracting anisotropically. The diffraction quality was poor and could not be improved despite major effort. Fab fragments were therefore generated and applied to co-crystallization with AgcS to improve crystal contacts (Supplementary Fig. 1a). Crystals of AgcS in a complex with 7B4 Fab fragment diffracted to ~3.2 Å. The phases obtained by a combination of molecular replacement, using a generic Fab structure, and experimental phasing from Se-Met labeled AgcS were of high quality, allowing us to unambiguously discern two AgcS molecules and two Fab fragments in one asymmetric unit (Supplementary Fig. 1b). In total, two structures of AgcS, one with D-alanine bound and the other with D-alanine bound, were determined (Supplementary Table 1).

AgcS forms a roughly cylindrical shape, which is ~40 Å tall and ~35 Å in diameter comprising 11 transmembrane (TM) helices connected by short cytoplasmic or periplasmic loops (Fig. 1, Supplementary Fig. 2a). Its N-terminus is at the periplasm while its C-terminus is cytoplasmic. AgcS appears to maintain pseudo 2-fold symmetry even though it contains an uneven number of TMs (Fig 1a, Supplementary Fig. 2b). One half contains TMs 2,3,7,6 and 8 while the second half contains TMs 4,5,9,10 and 11. TM1 resides outside of the helical bundle and does not seem to participate in translocation of substrates. The TMs vary greatly in their length and angle with relation to the plane of the membrane: some helices such as TM3 and TM9 appear almost perpendicular to the membrane plane while TMs 1 and 11 are heavily tilted. TMs 1, 3, 5, 6, 8, 10 and 11 surround the functional core of AgcS, which is made of TMs 2, 4, 7 and 9. Despite lack of sequence identity (15.1% to LeuT, 14.3% to ApcT and 14.8% to AdiC), the overall fold of AgcS is similar to that of LeuT (15, 21–23) from the NSS family. Superposing the crystal structures of AgcS and LeuT gives an overall root-mean-square deviation (RMSD) of 4.5 Å. The present structure of AgcS mirrors a common model of the LeuT fold Na^+^ and/or H^+^-coupled symporters(24, 25), including ApcT(14) and AdiC(26–28) of the APC family, BetP(29) of BCCT family, vSGLT(30, 31) of the SSS family, and Mhp1(32) of NCS1 family.

**Figure 1.**
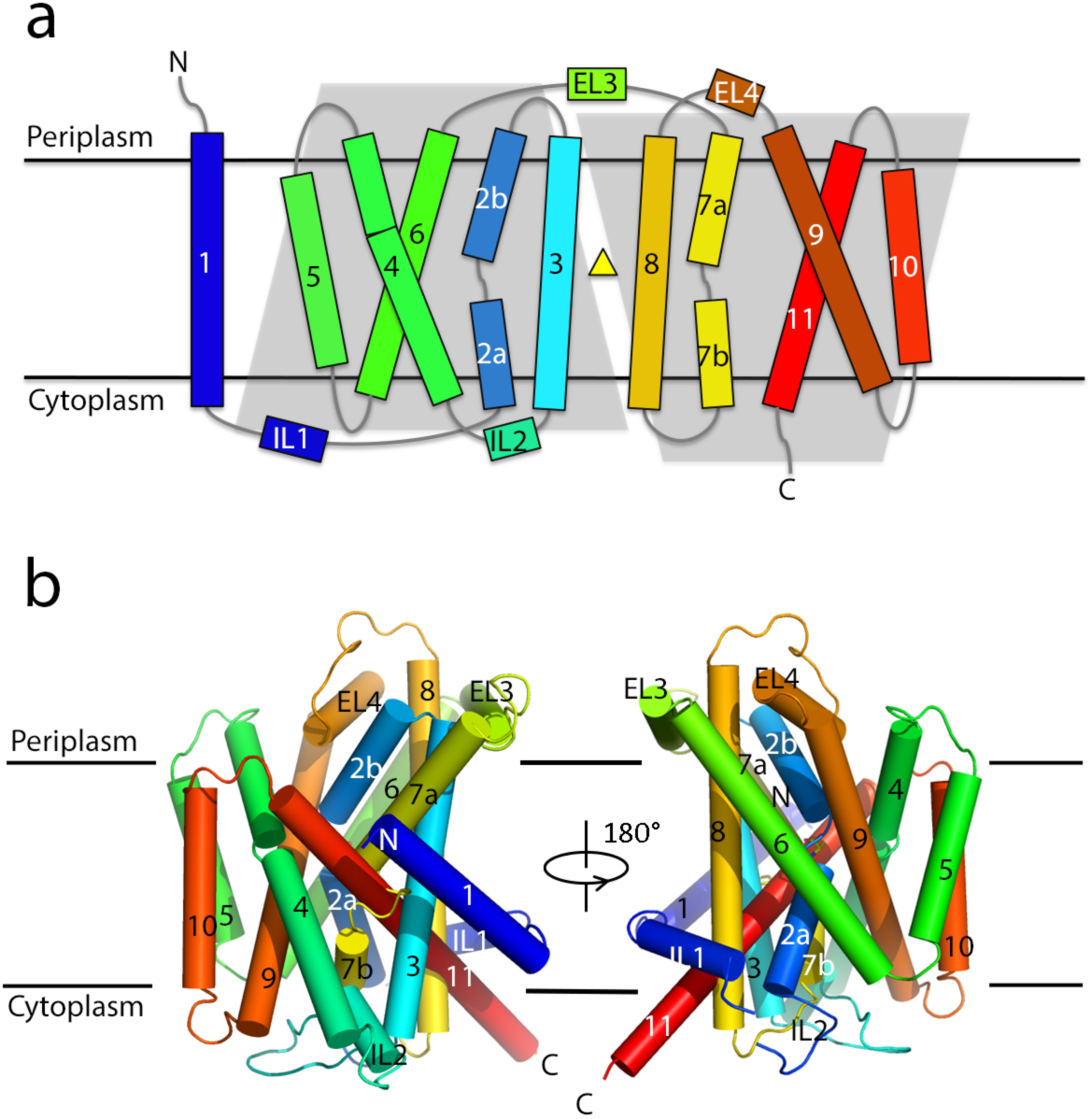
Overall structure of AgcS. (a). The topology of AgcS. The position of the bound alanine is depicted as a yellow triangle. (b). View in the plane of the membrane. Coloring and numbering of helices (cylinders) are the same as in Fig. 1A.

Notably, as with other LeuT-like symporters, two of the functional core helices in AgcS are kinked (Fig. 2 inset). TMs 2 and 7 line the substrate translocation pathway in AgcS and are broken into two segments. In TM2, Ile 76 and Gly 77 adopt an extended, non-helical conformation, linking segments 2a and 2b. TM7 contains a longer non-helical region from Glu 276 to Ser 281 connecting segments 7a and 7b. The residues at these non-helical loops expose carbonyl and amide groups at the center of the membrane bilayer for hydrogen bonding and ion coordination important for substrate binding and transport.

**Figure 2.**
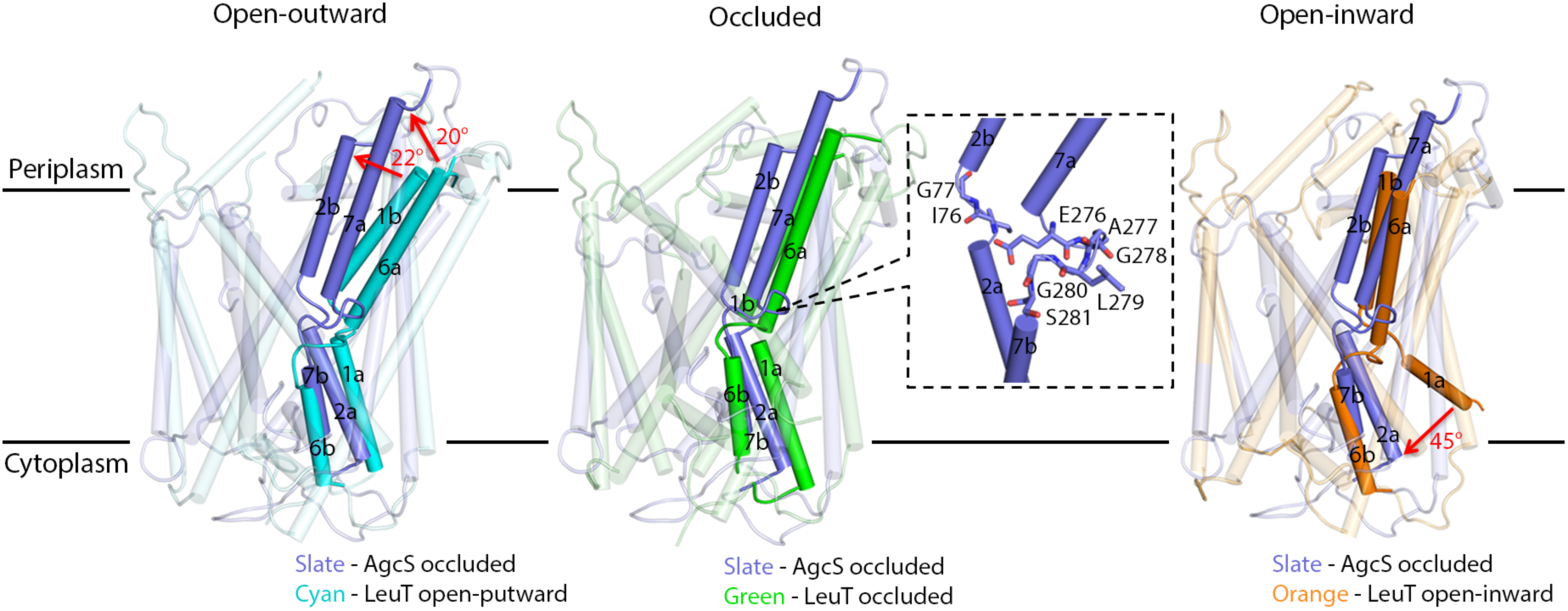
AgcS in the occluded state. AgcS structure was compared to LeuT open-outward state (left, PDB: 3tt1, cyans), LeuT occlude state (central, PDB: 2qei, green), and LeuT open-inward state (right, PDB: 3tt3, orange). Key structural changes are marked with red arrows. A close-up view of the unwound regions splitting TMs 2 and 7 are shown inset highlighting residues that are involved in substrate binding.

### AgcS structure in an occluded state

It is generally accepted that secondary transporters function though an alternate access model(33). The LeuT-fold transporters naturally alternate among three major conformations during their functional cycle: outward-open, occluded and inward-open state. In the outward-open state, a hydrophilic tunnel emerges and connects the substrate binding site located in the center of transmembrane helical bundle with the periplasm. When in the inward-open state, a hydrophilic tunnel leads from the substrate at the central binding site to the cytosol. During the occluded state, the substrate binding site is isolated from either the periplasm or cytoplasm. Within a complete switch cycle, substrates can be transported across the lipid membrane as the transporter alternates from one conformation to the others.

Two structures of AgcS were solved in this study, one with L-alanine and one with D-alanine bound, both captured in the occluded state (Fig. 2). No solvent accessible tunnels or vestibules were apparent leading from the substrate binding pocket across the membrane portion of the transporter. Previous studies reported the structures of LeuT in the open-outward, occluded and open-inward states(22, 23). Amino acid transporters such as AgcS and LeuT appear to gate through movements of the functional core helices. In AgcS these are TMs 2 and 7 while in LeuT they are TMs 1 and 6. As discussed above, these helices are broken (Fig. 2 inset) to allow free movement of the top half of the helices with respect to the bottom half and this in turn opens the binding pocket either to the outside or the inside of the cell. The pivot point appears to be at the substrate binding pocket. In the overlay of AgcS with open-outward LeuT (Fig. 2 left), or the overlay of AgcS with the open-inward LeuT (Fig. 2 right), core helices deviate by more than 20° or 45°, respectively. No such deviation was observed when occluded LeuT was overlaid with AgcS (Fig. 2 central). This analysis suggests that AgcS was captured in the occluded state.

A system of intricate amino acid interactions forms the extracellular and cytoplasmic gates of AgcS. The extracellular gate of AgcS consists of residues from TM2, TM4, and TM9 (Fig. 3a). Specifically, Thr 78 and Ala 73, on the two ends of the unwound region in TM2, expose their Cα nitrogen and carbonyl oxygen atoms for hydrogen bonding with Gln 170 (TM4) and Thr 366 (TM9). On the cytoplasmic side of the binding pocket, the main chain carbonyl oxygen atoms of Ala 74 (TM2) and one of the carboxylate oxygen atoms of Glu 276 (TM7) accept two additional hydrogen bonds from the indole nitrogen of Trp 370 (TM9), forming a plug-like motif to block the transport funnel. This hydrogen-bond network fastens TMs 2 and 7 to TMs 4 and 9 and lock the helices core bundle (Supplementary Fig. 2b) in an occluded state.

**Figure 3.**
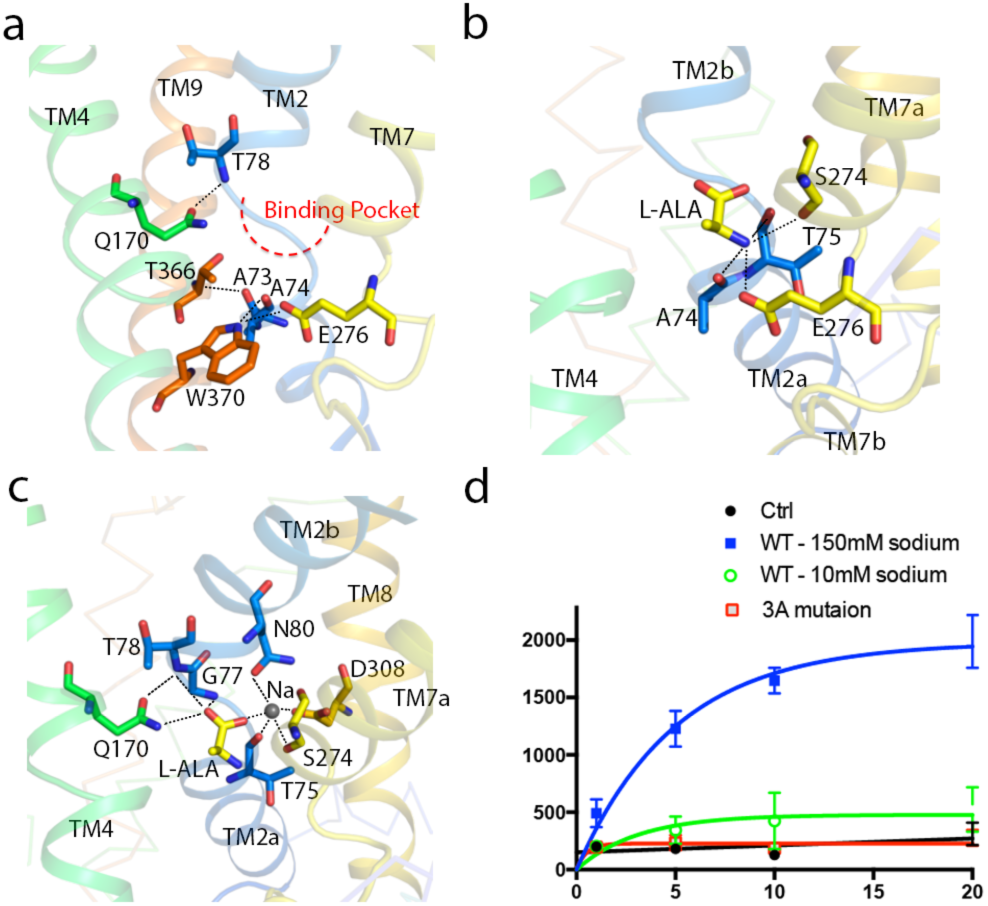
Gating interactions and substrate binding site in AgcS. (a). A close-up view of gating interaction network around binding pocket. (b). Hydrogen bonds between the amino group of the substrate and the residues of AgcS. (c). Hydrogen bonds and ionic interactions in the hydrophilic part of binding pocket are depicted as dashed lines. Bound Na^+^ ion shown as a grey sphere. (d). The relative uptake ability for missense mutant (three points made up of N80A, S274A, and D308A) compared with that of the WT protein.

### AgcS substrate binding and ion modulation

In current study, both structures of AgcS reveal a small amphipathic substrate binding cavity defined by residues from TMs 2, 4, 7 and 8. These residues define two distinct regions in the binding pocket: a hydrophilic charged region close to the extracellular gate and a hydrophobic region embedded deeply in the pocket toward the cytoplasmic gate. The hydrophobic pocket is mainly formed by Ile165 (TM4), Phe 273 (TM7), Thr 366 (TM9) and Trp 370 (TM9) (Fig. 4a).

**Figure 4.**
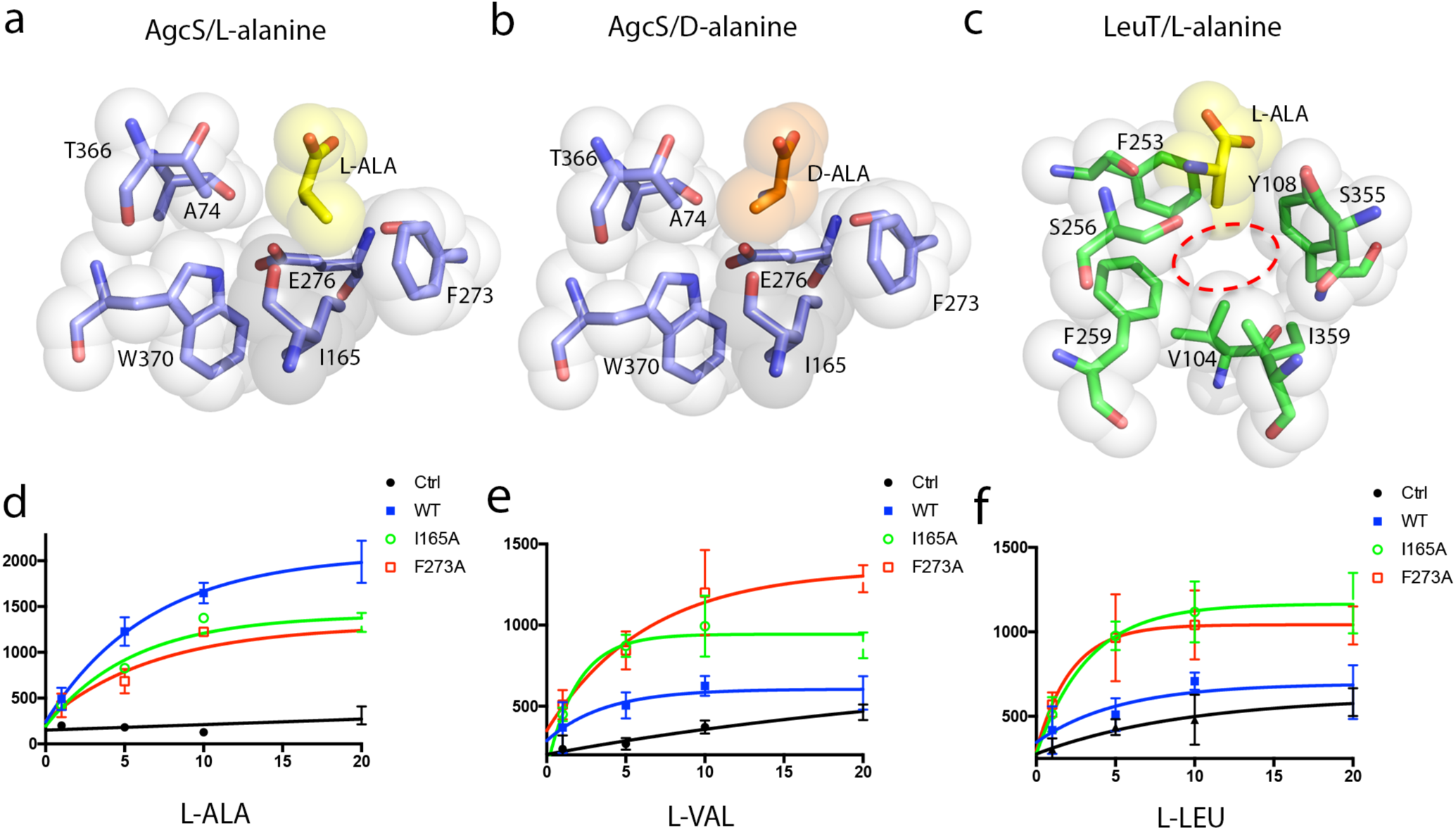
Substrate selectivity in AgcS dictated by the size of the binding pocket. Comparison of the hydrophobic pocket of alanine-bound AgcS and LeuT (PDB: 2qei). (a-c) Van derWaals surfaces for alanine side chain and interacting residues are shown as colored spheres. The binding pocket of AgcS is very small and only glycine or alanine can fit while in LeuT where substrate selectivity is broader the site is much larger able to accommodate larger amino acids (d-f) AgcS mutants I165A and F273A lose substrate selectivity allowing valine and leucine to be transported.

The hydrophilic part of the binding pocket contains both the amino and carboxyl groups of the bound alanine as well as a bound sodium ion. The amino group of the bound alanine is coordinated by the main-chain carbonyl oxygens of Ala 74, Thr 75 in TM2, and Ser 274 in TM7, and a side-chain carboxyl group from Glu 276 in TM7 (Fig. 3b). On the opposite side, the exposed main-chain of Gly 77 and Thr 78, which are in the unwound region of TM2, make contacts with the carboxyl group of alanine by direct hydrogen bonds, along with side-chain amide of the conserved Gln 170 (TM4) (Supplementary Fig. 3). This glutamine may also stabilize the irregular structure around the unwound region in TM2 by interaction between its side-chain carbonyl oxygen to the main-chain amino group of Thr 78 (Fig. 3c).

The carboxylic group of the bound alanine is further anchored to Asn 80 (TM2), Ser 274 (TM7) and Asp 308 (TM8) through the salt-bridge interactions mediated by a single sodium ion (Fig. 3c). These residues are highly conserved in the AgcS family (Supplementary Fig. 3). The coordination of sodium ion is provided by the side-chain carbonyl oxygens of Asn 80 and Asp 308 and the hydroxyl oxygen of Ser 274, further stabilized by main-chain carbonyl oxygen of Ser 274 and Thr 75 (Fig. 3c). To characterize the role of the conserved residues in sodium ion coordination, we generated a mutant (3A mutation including N80A, S274A and D308A) and evaluated the impact on substrate uptake using proteoliposomes. The mutant markedly reduced the uptake ability of L-alanine suggesting that the sodium ion plays a key role in facilitating substrate binding and uptake (Fig. 3d).

### Substrate selectivity in AgcS

AgcS shows a high selectivity for small amino acids like glycine and alanine with a dependency on sodium. We first evaluated AgcS binding affinities to several amino acids using Isothermal titration calorimetry (ITC). While heat exchange was identified for the binding of glycine, and both L- and D-alanine, other amino acids that were tested did not elicit a response (Supplementary Fig. 4). Uptake studies using reconstituted proteoliposomes likewise indicated impeccable specificity in AgcS for glycine, L-alanine and D-alanine in the presence of sodium (Supplementary Fig. 5, Fig. 3d).

The crystal structures of AgcS with L-alanine or D-alanine bound indicate that the substrate binding site is small and able to accommodate only small amino acids while maintaining an environment favorable to both alanine enantiomers. In contrast to the polar region of the bound alanine (discussed above), the methyl side chain is surrounded within a hydrophobic pocket mainly formed by the side chain of Ile 165 in TM4, Phe 273 in TM7, Thr 366 and Trp 370 in TM9 (Fig. 4a and 4b). That hydrophobic pocket could accommodate both alanine enantiomers equally well. The small size of this hydrophobic pocket, shaped by the van der Walls surface of these binding pocket residues, ensures that only glycine or alanine could fit while other amino acids not. In sharp contrast, the substrate binding pocket in LeuT is much larger (Fig. 4c) consistent with the broader selectivity of LeuT and its ability to accommodate a large variety of amino acids. In AgcS, the aliphatic side chains of any amino acid larger than alanine would clash with Ile 165 and Phe 273. Consistent with this postulate, mutation of I165A and F273A show poor substrate selectivity allowing additional amino acids such as L-valine and L-leucine to be efficiently transported by the mutated AgcS (Fig. 4d, 4e and 4f). These results indicate that the small size of the substrate binding pocket in AgcS is a key determinant in substrate selectivity, a structural feature that may apply to other APC superfamily members.

Stereo-selectivity and the unusual ability of AgcS to transport both enantiomers of alanine is of interest. Most amino acid transporters of the APC superfamily have effective stereo-selectivity and tend to transport L-type amino acids exclusively, mirroring the fact that L-amino acids are most commonly used in life while D-amino acids are rarely used. However, AgcS has an unusual ability to transport both enantiomers of alanine to be used as important nitrogen sources. This alanine utilization plays a crucial role to support *Methanococcus maripaludis* in harsh environments. As discussed above, the substrate binding site in AgcS has a cytoplasmic facing hydrophobic surface used to accommodate the aliphatic part of either L- or D-alanine equally well (Fig. 5a and 5b). In sharp contrast, transporters such as LeuT, that only allow L-type amino acids to be transported do not have an even distribution of hydrophobic surface at their substrate binding site. Instead, the cytoplasmic face of their substrate binding site is amphipathic containing both a hydrophobic patch and a charged polarized patch which is also spatially constrained (Fig. 5c). This feature ensures that only the L-enantiomers of amino acids can bind while D-enantiomers are repelled electrostatically.

**Figure 5.**
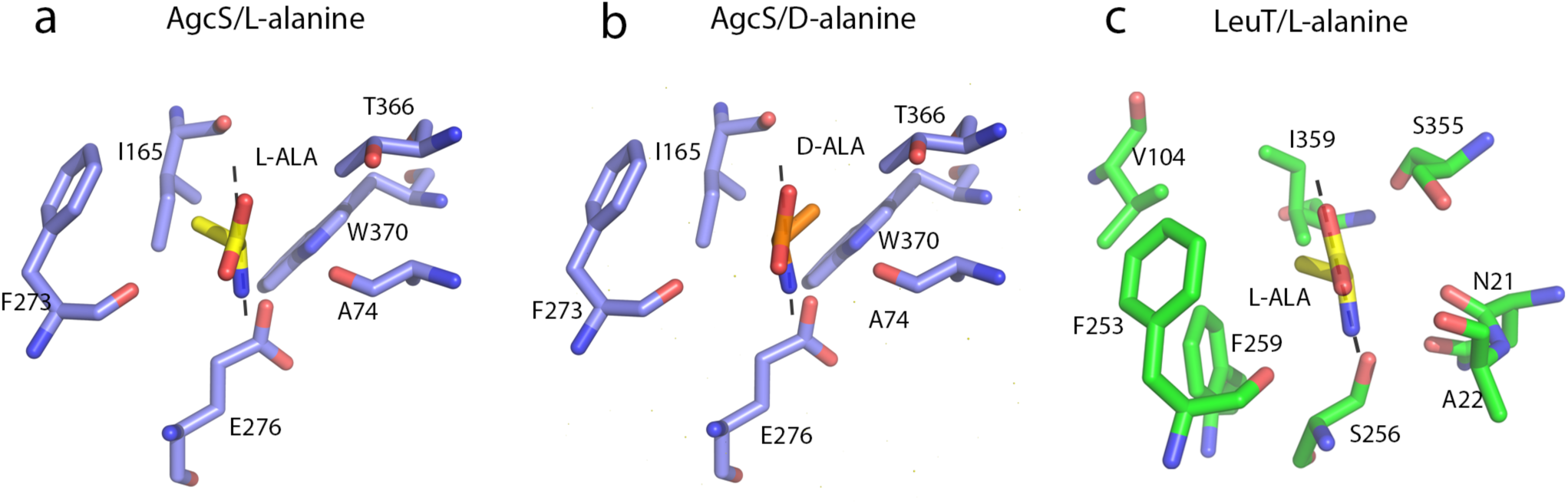
Electrostatics in the binding pocket of AgcS dictate stereoselectivity. Stereo-selectivity is afforded by electrostatic repulsion. The amide plane of substrate is shown as gray dash. In AgcS both alanine enantiomers fit equally well as the two faces of the binding pocket are hydrophobic. In LeuT, which only binds L-amino acids, one face is hydrophobic while the other is charged ensuring selection for L-amino acids over D-amino acids.

### Concluding remarks

The mechanism of substrate binding and selectivity is an essential feature for the APC superfamily members. Here we described the crystal structure of an alanine transporter AgcS capable of transporting both L- and D-alanine and discovered that both the size and polarity of the substrate binding pocket are important structural determinants for specificity and stereo-selectivity. The size of the binding pocket determines what amino acids could fit in the transporter while electrostatics control whether L-type or D-type amino acids could bind. We showed that a single point mutation in the binding site of AgcS can broaden the specificity of this transporter to allow uptake of larger amino acids such as valine and leucine. This study forms the basis for future studies on stereo-selectivity in nature and for the rational design of new transporters.

## METHODS SUMMARY

AgcS (Gene ID: 2761075) from *Methanococcus maripaludis* and its mutants were overexpressed in E. coli BL21 (DE3) C43. Fab antibody fragments were generated as described in Methods. The AgcS-Fab complex was purified in the presence of 0.2% (w/v) n-decyl-β-D-maltoside and crystallized in the following condition, 0.1 M Sodium Citrate pH 5.6, 2.5 M Ammonium Sulfate. Crystals were obtained in the presence of 50 mM L-alanine and D-alanine. Diffraction data sets of all crystals were collected at the Advanced Light Source (Beamline 8.2.1 and 5.0.2) and Advanced Photon Source (NE-CAT 24-ID-C). Data processing and structure determination were performed using the autoPROC, RAPD, PHENIX and COOT programs. Detailed methods can be found in the supplementary information section that accompany this manuscript.

## Acknowledgements

We thank D. Cawley for development and production of monoclonal antibodies, and staff at the Berkeley Center for Structural Biology, Advanced Light Source (ALS) and the Northeastern Collaborative Access Team (NE-CAT), located at the Advanced Photon Source (APS), for assistance with X-ray data collection. The Berkeley Center for Structural Biology is supported in part by the National Institutes of Health, National Institute of General Medical Sciences, and the Howard Hughes Medical Institute. The Advanced Light Source is a Department of Energy Office of Science User Facility under Contract No. DE-AC02-05CH11231. This work is also based upon research conducted at the Northeastern Collaborative Access Team beamlines, which are funded by the National Institute of General Medical Sciences from the National Institutes of Health (P41 GM103403). The Pilatus 6M detector on 24-ID-C beam line is funded by a NIH-ORIP HEI grant (S10 RR029205). This research used resources of the Advanced Photon Source, a U.S. Department of Energy (DOE) Office of Science User Facility operated for the DOE Office of Science by Argonne National Laboratory under Contract No. DE-AC02-06CH11357. Research in the Gonen laboratory is funded by the Howard Hughes Medical Institute. Coordinates and structure factors were deposited in the Protein Data Bank.

